# Divisive Normalization Circuits Faithfully Represent Auditory and Visual Sensory Stimuli

**DOI:** 10.1101/2022.09.17.506431

**Authors:** Aurel A. Lazar, Tingkai Liu, Yiyin Zhou

## Abstract

Divisive normalization is a canonical neural computation employed for sensory adaptation in vision, olfaction and attention modulation. While Divisive Normalization has been proposed to be an efficient coding algorithm, it remains unclear whether such transformation results in information loss in a dynamic setting. Leveraging a previously proposed general mathematical framework called the Divisive Normalization Processor (DNP), we first show that the DNP circuit describes a wide class of neural circuits as well as phenomenological models including the Linear-Nonlinear cascade model. We then demonstrate both theoretically and computationally that the DNP is an invertible operator that could faithfully represent input information given sufficient output samples.

## Executive Summary

### Overall Message

Given a *sufficient* number of samples, the Divisive Normalization Processor operating on audio scenes and visual fields is invertible.

### Problem Setting

Divisive normalization underlies many models of neural circuits arising in auditory and visual systems. Due to the highly nonlinear nature of the divisive normalization operation in a number of applications such as contrast gain control, there has been a lack of quantitative characterization of the information content of the signals at the output of these circuits. In this paper, we show that, stimuli processed by a class of circuits that we call divisive normalization processors can be faithfully recovered from the outputs of the circuit, given a sufficient number of samples. We demonstrate this for auditory stimuli (Figure 1A) and visual stimuli (Figure 1B).

**Figure 1:**
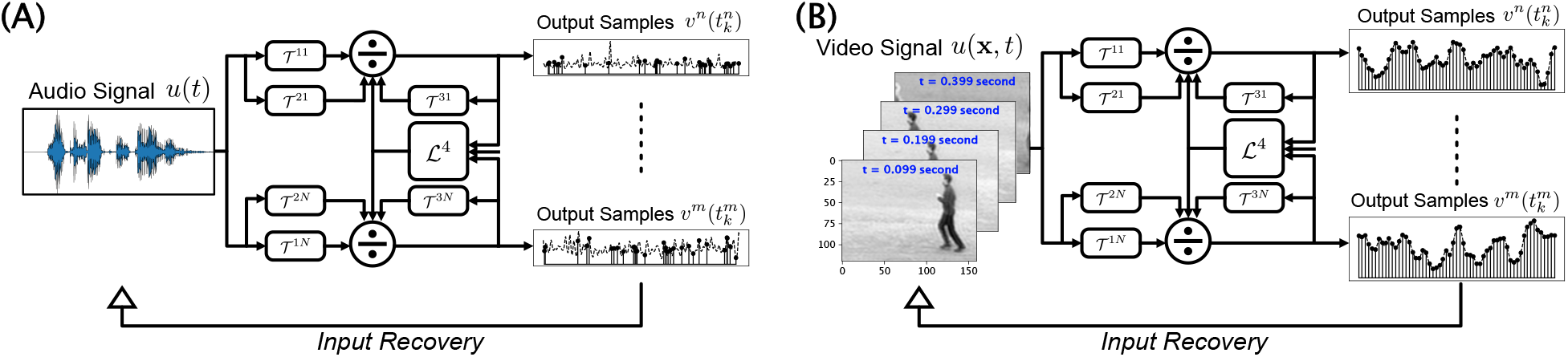
(A) Divisive Normalization Processor (DNP) encoding and input recovery of auditory stimuli. Note that the output signals are asynchronously sampled. (B) DNP encoding and input recovery of video stimuli.

### Context/Relevance to the Published Literature

- DNP processing underlies many existing models of auditory systems, e.g., [1].
- For a model of the inner ear with gammatone filterbank and synthetically generated Volterra kernels, audio stimuli can be faithfully recovered from the outputs of the DNP.
- DNP processing underlies many existing models of visual circuits, for example, [2] and [3].

### Formal Modeling of DNPs

1. We present the modeling of auditory and visual stimuli.
2. We introduce feedforward and feedback DNPs as depicted in Figure 2, and formally describe the I/O of the DNPs.

**Figure 2:**
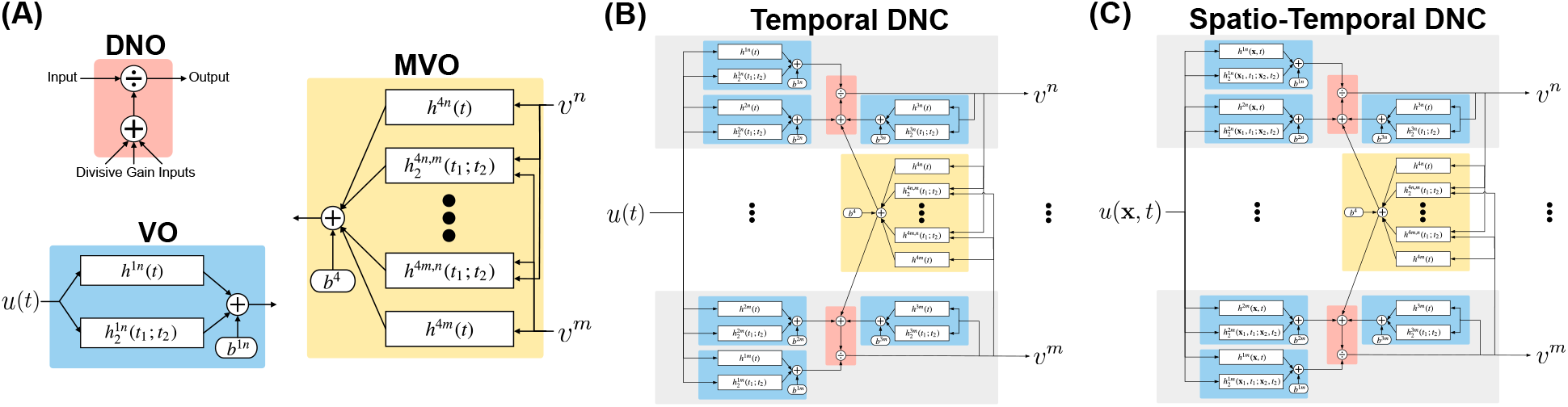
Divisive Normalization Processors. (A) Components of DNPs. (B) DNP for temporal input signal *u*(*t*). (C) DNP for spatio-temporal input signal *u*(**x**, *t*).

### Formulating the Recovery of Input Stimuli and Evaluation Criteria

#### Formulating the Recovery of Input Stimuli

Given the input/output relationship of the DNP as

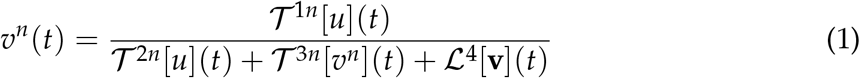

and irregularly sampled output values *v^n^*(*t_k_*),*n* = 1,…,*N,k* = 1,…,*N_K_*, the recovery problem is formulated as finding *u* given the samples *v^n^* (*t_k_*) with known operators 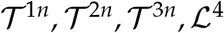.

#### Evaluation Criteria

Given the recovered input signal *ũ*, we evaluate the recovery via Signal-to-Noise Ratio (SNR) as well as Peak Signal-to-Noise Ratio (PSNR):

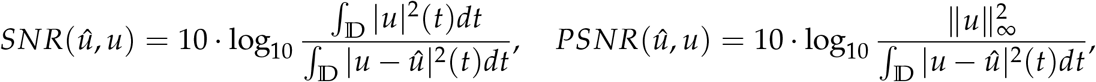

where *u* = *u*(*t*) is the temporal stimulus, *û* = *û*(*t*) is the recovered stimulus. For spatio-temporal stimuli, *u* = *u*(**x**, *t*) is the stimulus and *û* = *û* (**x**, *t*) is the recovery. 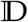 defines the domain of the stimuli.

### Perfect Recovery Algorithms

#### Stimuli Processed by DNPs with First Order Kernels Are Recovered via Regularized Regression

- Sampling of the DNP output can be asynchronous (this is a processor with continuous input and continuous output)
- The reconstructed stimulus is the solution of the *convex* optimization problem

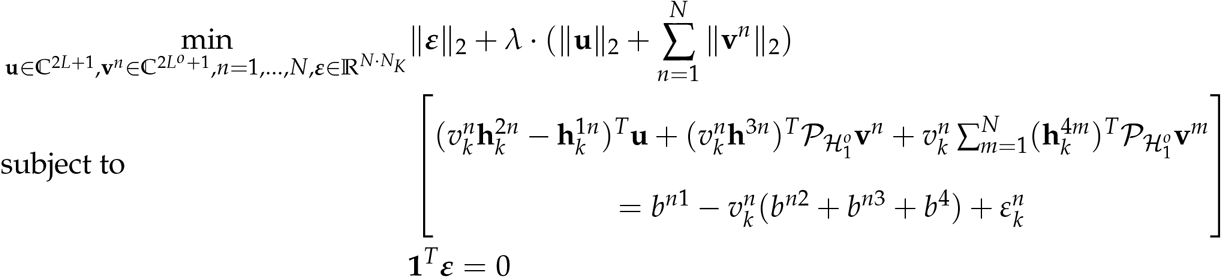

where the expression in [·] brackets is the I/O constraint of the DNP with 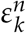 denoting the error of measurement at the *k*-th sample of the *n*-th neuron.
- Without feedback, the total number of samples required is 2*L* + 1 (same as the order of the input signal), with feedback the total number of samples required is 2*L* + 1 + *N*(2*L*^o^ + 1) (proportional to the number of channels). The extra samples are needed to recover the outputs from the samples according to the Shannon-Nyquist sampling theorem.

#### Stimuli Processed by DNPs w/ Second Order Kernels Are Recovered via Rank-Minimization

- With 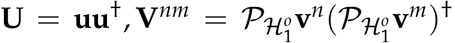, where ^†^ denotes the conjugate transpose, and by observing that **U** and **V**^*nm*^ are rank 1, we formulate the recovery problem as a rank-minimization problem with a nuclear-norm *convex* relaxation:

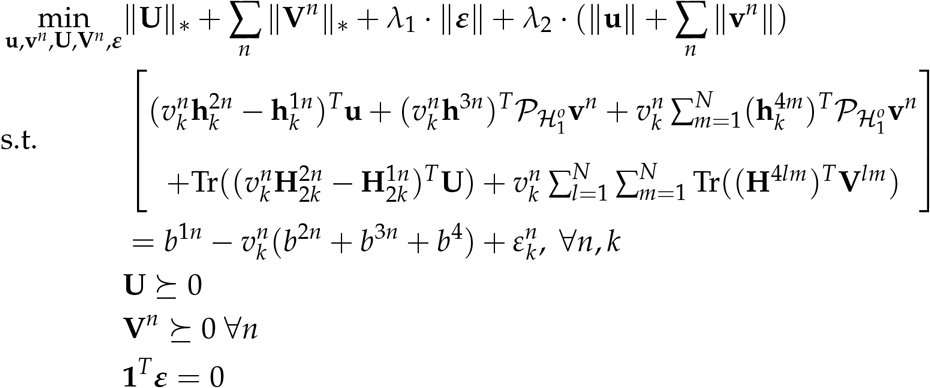

where the first constraint above is the I/O of DNP with first and second order kernels, 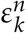. is the error of the measurement at the *k*-th sample of the n-th neuron, ||·||_*_ denotes the nuclear norm operator, and *λ*_1_, *λ*_2_ ≥ 0.
- Given a sufficient number of samples, the recovery is perfect and the recovered 2nd order coefficient is rank-1.

We use simple temporal stimuli to characterize how well the stimulus is recovered with an increasing number of samples. Perfect recovery can always be achieved when the number of samples is above a threshold (Figure 3).

**Figure 3:**
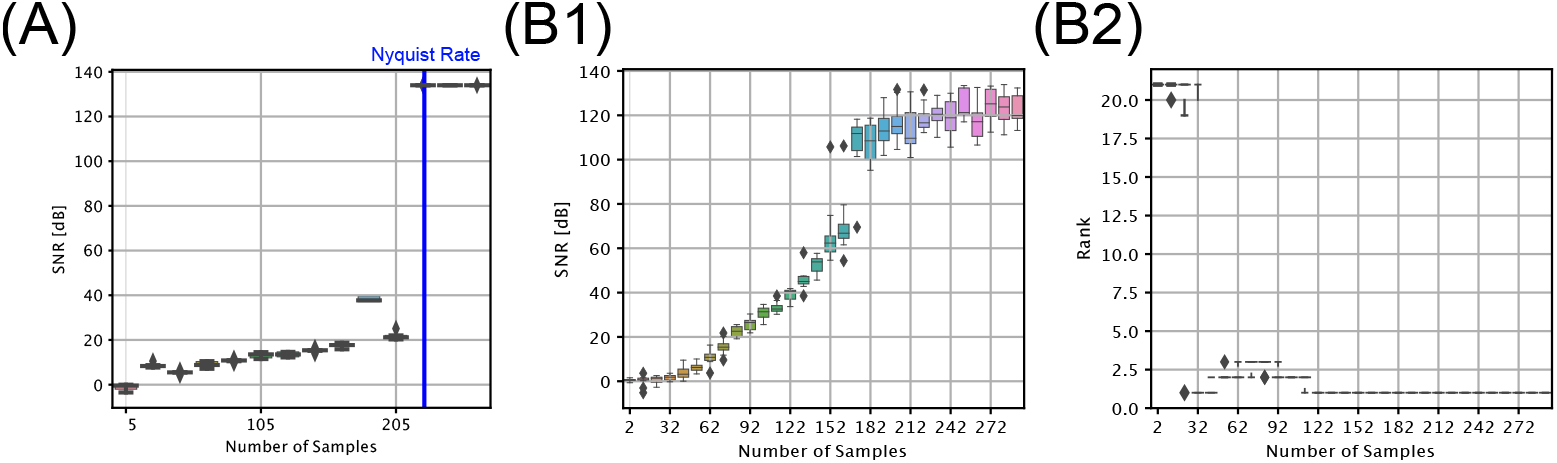
Input recovery for DNP circuit with 1st (A) and 2nd (B1,B2) order kernels. (A,B1) Given a sufficient number of samples, the input recovery for DNP circuit reaches Signal-to-Noise ratio above 133 dB, indicating perfect recovery. (B2) For DNP circuit with 2nd order kernels, perfect input recovery correponds to when the recovered 2nd order input coefficient **U** has rank 1.

## Acknowledgments

The research reported here was supported by AFOSR under grant #FA9550-16-1-0410, DARPA under contract #HR0011-19-9-0035 and NSF under grant #2024607.

